# Oxytocin alleviated colitis and colitis-associated colorectal tumorigenesis by targeting fucosylated MUC2

**DOI:** 10.1101/2023.08.17.553684

**Authors:** Xia Wang, Dawei chen, MengNan guo, Yao Ning, Jing Guo, Jiahui Gao, Xiaoran Xie, Dong Zhao, Lixiang Li, Shiyang Li, Yanqing Li, Xiuli Zuo, Jingxin Li

**Author notes:** Correspondence: X. Z. or J. L.

## Abstract

Colon cancer is commonly regarded as hormone-independent. However, there have been reports suggesting the involvement of sex hormones in colon cancer development. Nevertheless, the role of hormones from the hypothalamus-hypophysis axis in colitis-associated colorectal cancer (CAC) remains uncertain. In this study, we observed a significant reduction in the expression of the oxytocin receptor (OXTR) in colon samples from both colitis and CAC patients. To investigate further, we generated mice with an intestinal epithelium cell (IEC)-specific knockout of OXTR. These mice exhibited markedly increased susceptibility to dextran sulfate sodium (DSS)-induced colitis and DSS/Azoxymethane (AOM)-induced CAC compared to wild-type mice. Our findings indicate that OXTR depletion impaired the inner mucus of the colon epithelium. Mechanistically, oxytocin was found to regulate MUC2 maturation through B3GNT7-mediated fucosylation. Interestingly, we observed a positive correlation between B3GNT7 expression and OXTR expression in human colitis and CAC colon samples. Moreover, the administration of oxytocin significantly alleviated tumor burden. Hence, our study unveils oxytocin’s promising potential as an affordable and effective therapeutic intervention for individuals affected by colitis and CAC.

## Introduction

Inflammatory bowel disease (IBD), consisting of ulcerative colitis (UC) and Crohn’s disease (CD), poses a significant global public health challenge (1). These conditions lead to debilitating gastrointestinal symptoms and progressive inflammation, resulting in irreversible damage to the gastrointestinal tract and an escalated risk of CAC. The incidence of CAC is approximately 20% in UC and 8% in CD (2). Therefore, understanding the pathogenesis of IBD and its association with CAC has important implications for the management of IBD patients.

The mucus layer, acting as the primary barrier between intestinal bacteria and intestinal epithelial cells (IECs) (3), plays a critical role in the pathogenesis of IBD (4). Mucin2 (MUC2), synthesized by intestinal goblet cells, is a key component of the mucus layer and crucial for colonic protection (5). However, the molecular mechanisms underlying the role of MUC2 in CAC remain unclear.

Oxytocin (OXT), a neuro-hypophysial hormone and neuropeptide primarily synthesized by the hypothalamus and released by the pituitary gland,(6) exerts its effects through the oxytocin receptor (OXTR). OXTR belongs to the G protein-coupled receptor (GPCR) class A/rhodopsin family and plays a pivotal role in social bonding, stress response, maternal behavior, sexual activity, uterus contraction, milk ejection, and cancer (7). Emerging evidence suggests the presence of both OXT and OXTR in the gastrointestinal system (8), implicating them in gastrointestinal tumorigenesis (9, 10). Studies, including our own, have demonstrated the potential of OXT to alleviate experimental colitis (11) and modulate the anti-inflammatory response (12). Notably, the deletion of OXT signaling in macrophages (13) and dendritic cells exacerbates DSS-induced colitis (14). However, the role of OXT/OXTR in IECs remains poorly understood, necessitating further mechanistic investigations in gastrointestinal diseases.

This study aimed to uncover the essential function of OXT signaling in colonic carcinogenesis and colitis. We hypothesized that the deficiency of OXTR in IECs may render them more susceptible to CAC and UC. Mechanistically, we investigated the impact of OXT/OTR on the mucosal barrier function by examining the transcriptional activation of β1-3-N-acetylglucosaminyltransferase7 (B3gnt7) and the induction of α1-3-fucosylation of MUC2. Our findings provide valuable insights into a novel diagnostic indicator for CAC and colitis, contributing to a better understanding and management of these conditions.

## Results

### IECs Ablation of OXTR Sensitizes Mice to CAC

The dysregulation of hormones is frequently associated with cancer initiation and progression (15). The hypothalamus-hypophysis axis, which controls the production and secretion of hormones through the pituitary portal vein system, regulates various physiological functions of the body (16). Here, we examined the expression of relevant receptors in colitis-associated cancer (CAC) using the GEO database and found that the expression of GHRHR and OXTR genes was decreased (Figure. 1A). Since the role of GHRHR in CAC is well-established (17), we focused on oxytocin. We observed decreased expression of OXTR in CAC patients compared to adjacent healthy tissue (Figure.1, B and C). Consistent with these findings, mRNA expression of OXTR was also weakened in a mouse colon cancer model induced by AOM/DSS (Figure. 1D). To examine the functional role of OXTR in CAC, we established an AOM/DSS model in OXTR^fl/fl^ and OXTR^△IEC^ mice (Figure. 1E). Following the AOM/DSS protocol, OXTR^△IEC^ mice exhibited significantly increased tumor burden compared to OXTR^fl/fl^ mice (Figure. 1F). Statistical analysis revealed that OXTR^△IEC^ mice had a shorter colon length (Figure. 1G), a higher number of tumors (Figure. 1H), a wider tumor area and tumor load (Figure. 1I), and increased spleen weight (Figure. 1J). RT-qPCR analysis showed increased expression of inflammatory factors (Figure. 1K). H&E staining demonstrated substantial tumor areas, more colon injury, and immune cell infiltration in the colon tissues of OXTR^△IEC^ (Figure. 1L). Moreover, the survival time of OXTR^△IEC^ mice was significantly shorter (Figure. 1M). Overall, these results indicate that OXTR deficiency in intestinal epithelial cells exacerbates AOM/DSS-induced CAC progression.

**Figure. 1.**
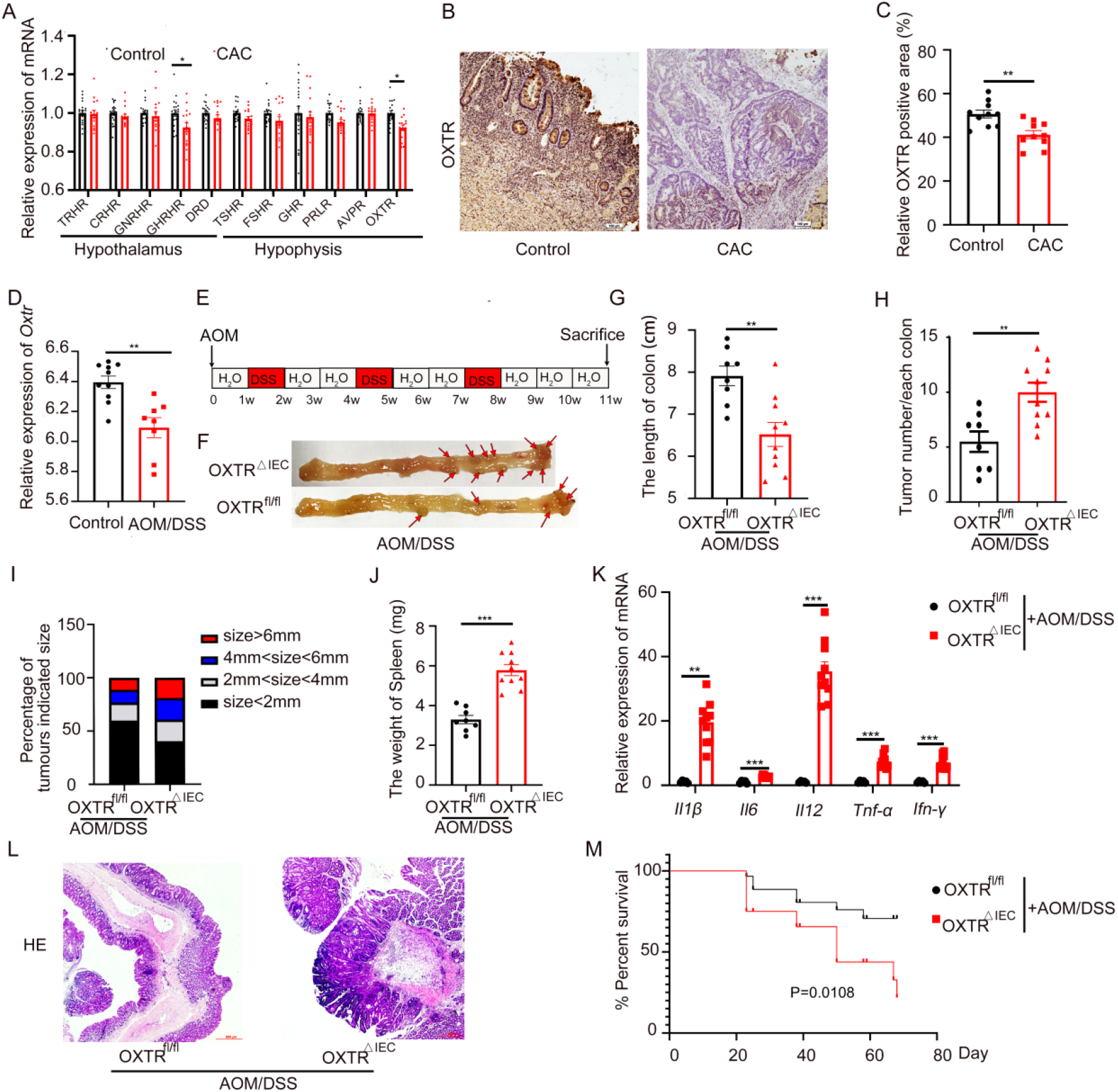
Ablation of OXTR in IECs promotes the development and progression of colitis-associated cancer. **(A)** Analysis of hypothalamus-hypophysis hormone receptors gene expression in normal and CAC patients. TRHR, thyrotropin-releasing hormone receptor; CRHR, thyrotropin-releasing hormone receptor; GNRHR, gonadotropin-releasing hormone receptor; GHRHR, growth hormone releasing hormone receptor; DRD, dopamine receptor D; TSHR, thyroid stimulating hormone receptor; FSHR, thyroid stimulating hormone receptor; GHR, growth hormone receptor; PRLR, prolactin receptor; AVPR, arginine vasopressin receptor; OXTR, oxytocin receptor, (GSE37283 and GSE47908). **(B)** Representative images of OXTR IHC staining of human normal and CAC tissue. Scale bar, 100 μm. **(C)** The quantifications of OXTR-positive area. N=5. **(D)** OXTR gene expression in AOM/DSS-induced colon cancer model. (GSE113002). E-M) OXTR^fl/fl^ (n=8) and OXTR^△IEC^ (n=10) mice were treated with AOM and 3 cycles of 2% DSS to induce inflammation-driven colorectal cancer. **(E)** Flow chart of the induction procedure. **(F)** Representative photos of the colons. **(G)** Colon length. **(H)** Number of tumors in the colon per mouse. **(I)** The percentage of different tumor diameters. **(J)** Spleen weight. **(K)** Relative inflammatory cytokine mRNA levels in the colon were measured by real-time polymerase chain reaction. **(L)** Representative H&E-stained colon sections. Scale bar, 500 μm. **(M)** Survival curve assessed by log-rank (Mantel-Cox) test. Data are presented as the mean ± SEM. *P < 0.5, **P < 0.1, ***P < 0.01. P-values were determined by using A, K) two-way analysis of variance (ANOVA) or C, D, G, H, J) unpaired two-tailed Student’s t-test.

## OXTR Deficiency in IEC Facilitates CAC Depends on Inflammation

The AOM/DSS-mediated primary cancer model involves synergistic effects of AOM-induced mutation and DSS-induced inflammation in CAC development (18). To determine whether AOM or inflammation is necessary for OXTR to suppress CAC, we evaluated a colitis-independent colon cancer model (Figure. S1A). Our results showed no significant difference in tumor incidence including colon length, tumor burden, spleen weight, histological characteristics, or inflammation levels between the two groups (Figure. S1, B-F). Additionally, we administered 2% DSS to OXTR^fl/fl^ and OXTR^△IEC^ mice to induce chronic colitis (Figure. 2, A-H). Compared with OXTR^fl/fl^ mice, OXTR^△IEC^ mice exhibited exacerbated colitis, including increased weight loss, disease activity index, and spleen weight, as well as enlarged histologic mucosal damage and higher levels of inflammatory cytokines under DSS-treated condition (Figure. 2, A-H). We also examined the role of OXTR in an acute colitis model, where mice were treated with 2.5% DSS for 6 days. Consistent with chronic colitis, OXTR^△IEC^ mice developed severe colitis, with more significant body weight reduction, higher stool score and bleeding scores, shorter colon length, larger spleen size, and increased inflammatory factors compared to OXTR^fl/fl^ mice (Figure. 2, I-N). These findings indicate that the suppressive role of oxytocin in colon cancer development is inflammation-dependent.

**Figure. 2.**
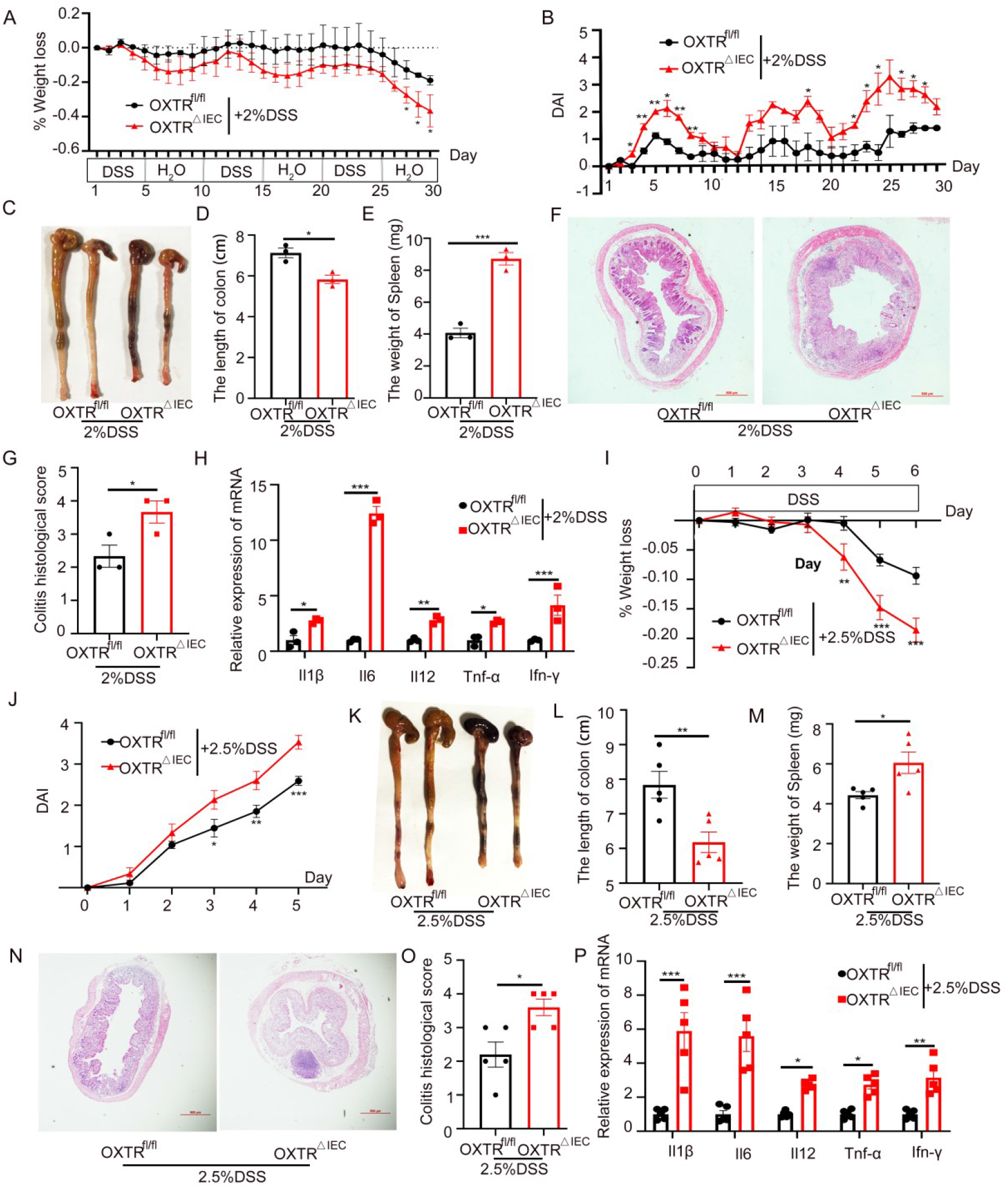
OXTR IEC-deficient mice develop severe DSS-induced colitis. **(A-H)** OXTR^fl/fl^ (n=3) and OXTR^△IEC^ (n=3) mice were treated with 2% DSS to induce chronic colitis. **(A)** Body weight change was shown as a percentage of initial body weight. **(B)** The disease activity index (DAI) was scored. **(C)** Representative photos of the colons. **(D)** Colon length. **(E)** Spleen weight. **(F)** Representative H&E-stained colon sections. Scale bar, 500 μm. **(G)** Semi-quantitative scoring of histopathology in colon tissue. **(H)** Cytokine production in whole cells from the colon. I-P) OXTR^fl/fl^ (n=5) and OXTR^△IEC^ (n=5) mice were treated with 2.5% DSS to induce severe colitis. **(I)** Body weights were measured daily and are depicted as a percentage of initial body weight. **(J)** DAI score. **(K)** Representative photos of the colons. **(L)** Statistical analysis of colon length. **(M)** Statistical analysis of spleen weight. **(N)** Representative H&E-stained images, scale bar, 500 μm. **(O)** Histologic injury score. **(P)** RT-qPCR analysis of the relative expression of inflammatory factors in colon tissues. Data are presented as the mean ± SEM. *P < 0.5, **P < 0.1, ***P < 0.01. P-values were determined by using A, B, H, I, J, P) two-way analysis of variance (ANOVA) or D, E, G, L, M, O) unpaired two-tailed Student’s t-test.

### IEC-Specific Ablation of OXTR Impairs the Integrity of the Inner Mucus Layer

To understand how OXT signaling modulates the pathogenesis of DSS-induced colitis, we evaluated the expression levels of colonic barrier proteins in OXTR^fl/fl^ and OXTR^△IEC^ IECs with or without DSS treatment. We found that the absence of OXTR did not affect the level of tight junction proteins but did impact the protein levels of the mucin MUC2, the dominant component of the inner mucus layer (Figure. S2 and Figure. 3, A-C). As the inner mucus layer, a physical barrier, prevents excessive inflammation by separating microbes, we examined its integrity (19). Our results showed that exposure to DSS led to a considerable decline in the thickness of the inner mucus layer (Figure. 3, D and E). Importantly, compared to OXTR^fl/fl^, the inner mucus layer was extensively damaged in OXTR^△IEC^ mice even under physiological conditions, and near-complete disappearance in DSS-treated OXTR^△IEC^ mice (Figure. 3, D and E). The inner mucus layer is primarily produced by colonic goblet cells (GCs), specialized cells located in the epithelium.(4) Interestingly, we found that the differentiation of GCs was comparable between OXTR-deficient and OXTR^fl/fl^ mice, although the expression of MUC2 was decreased (Figure. 3, F and G and Figure. S2, G and H). Notably, OXT upregulated MUC2 expression in LS174T cells and OXTR^fl/fl^ colonic organoids, while the inhibition of OXTR with L-368,899 abolished the effect of OXT. Furthermore, in OXTR-deficient organoids, OXT had no impact (Figure. S3, A-C). Collectively, these findings suggest that OXT/OXTR regulates the integrity of the inner mucus layer by modulating the expression of MUC2.

**Figure. 3.**
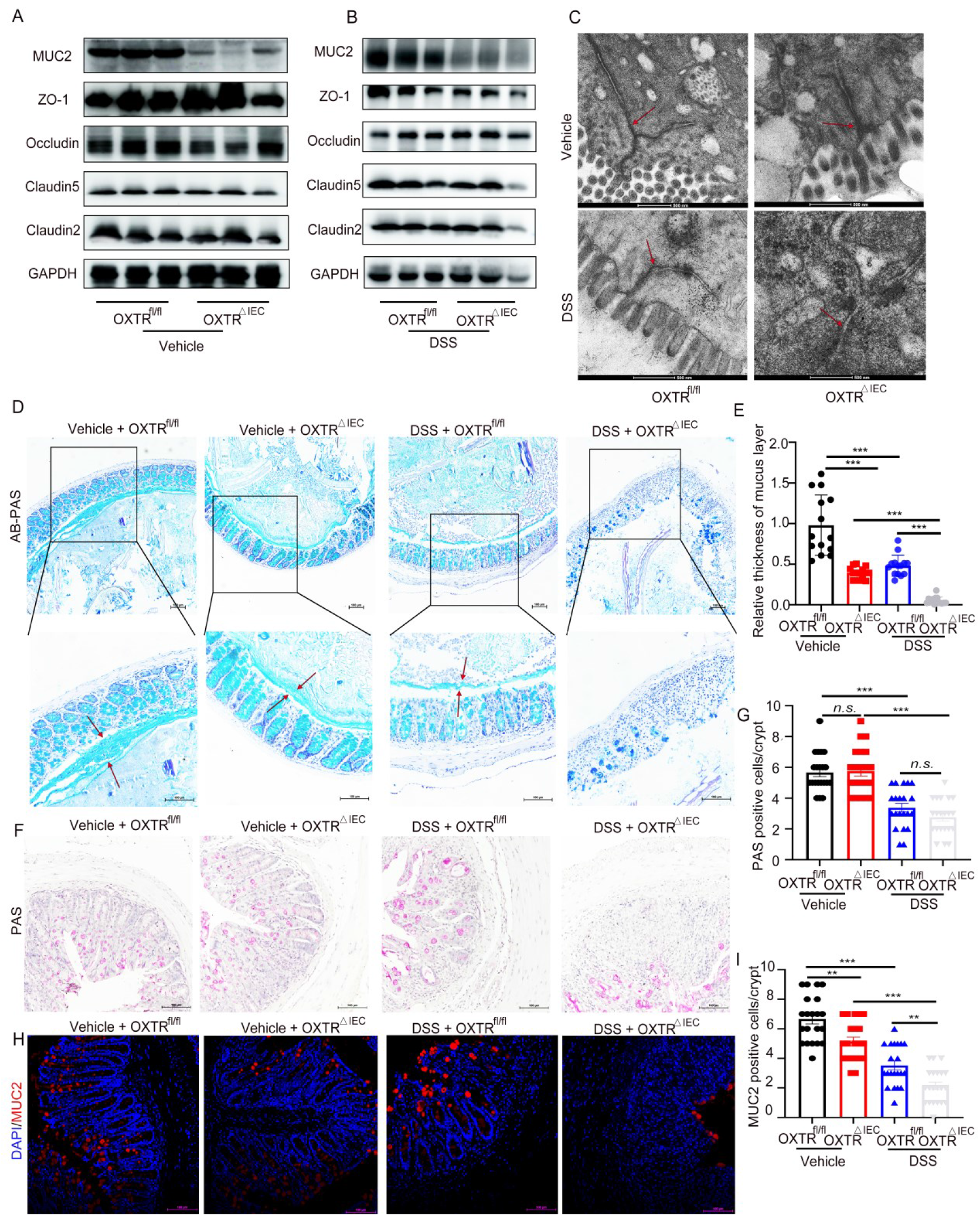
OXTR is important for preserving inner mucus layer integrity following challenges with DSS. (A-**B)** Western blot analysis of the barrier proteins level in the IECs from OXTR^fl/fl^ and OXTR^△IEC^ mice treated with or without 2.5% DSS. **(C)** Transmission electron microscopy of colons obtained from untreated mice or mice treated with 2.5% DSS. Scale bar, 500 nm. (n=3) **(D)** Representative micrographs of periodic acid-Schiff (PAS)-AB staining. Scale bar, 100 μm (n=3) **(E)** Quantifications of mucus thickness in colon sections with contents from vehicle-and DSS-treated mice. (n=3) **(F)** Representative micrographs of periodic acid-Schiff (PAS) staining. Scale bar, 100 μm. (n=3) **(G)** The number of PAS-positive cells was counted randomly in 7 crypts for each mouse. Histogram demonstrating the number of Goblet cells per crypt. (n=3). **(H)** Immunofluorescence staining of Muc2 (red) and nuclei (DAPI, blue), scale bar, 100 μm. (n=3) **(I)** The number of MUC2-positive cells was counted randomly in 5 crypts for each mouse. (n=3) Data are presented as the Mean ± SEM. n.s, no significant, **P < 0.1, ***P < 0.01. P-values were determined by using a two-way analysis of variance (ANOVA).

### Oxytocin Regulates MUC2 Expression via B3GNT7

To investigate the effect of OXT signaling on MUC2 expression, we conducted RNA sequencing (RNA-seq) analysis on OXTR^fl/fl^ and OXTR^△IEC^ IECs under physiological and pathological conditions. Differential gene expression analysis revealed several downregulated genes in both OXTR^fl/fl^ and OXTR^△IEC^ mice treated with or without DSS. Among these genes, B3gnt7, which is involved in the O-linked glycosylation of mucins, showed consistent downregulation in all groups (Figure. 4, A-D). Reverse transcription-polymerase chain reaction (RT-qPCR) and western blot analysis confirmed the decreased expression of B3GNT7 in OXTR^△IEC^ colonic epithelial cells (Figure. 4, E-H). Additionally, incubation with OXT upregulated B3GNT7 expression in LS174T cells, while treatment with the OXTR inhibitor L-368,899 inhibited the effect of OXT. Similar results were observed in colonic organoids lacking OXTR (Figure. S3, D-G). Moreover, downregulation of B3GNT7 using siRNA in LS174T cells and colonic organoids inhibited the OXT-dependent increase in MUC2 expression (Figure. 4, I and J). These findings demonstrate that the expression of MUC2 is regulated by the OXT/OXTR-B3GNT7 pathway.

**Figure. 4.**
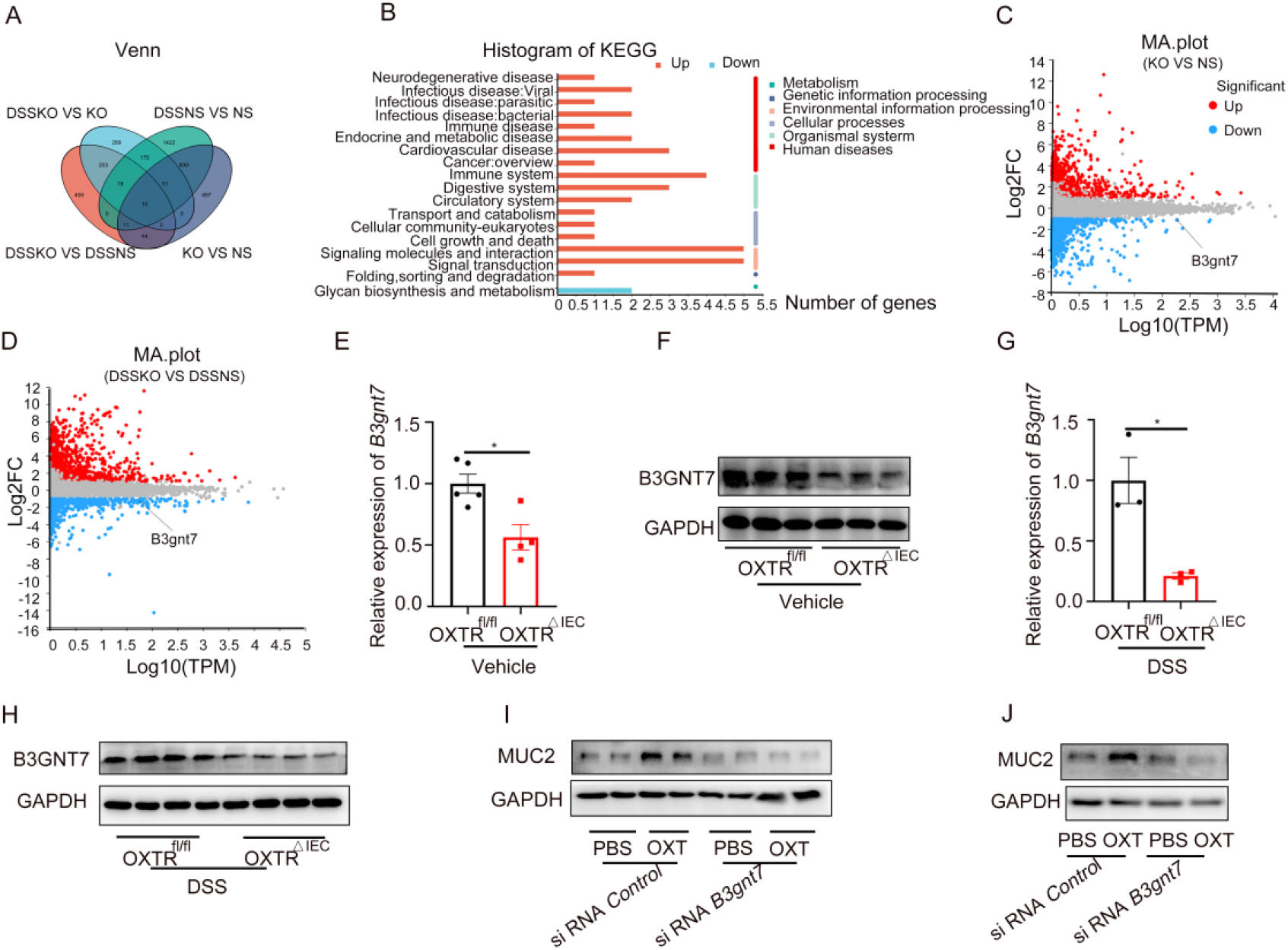
Oxytocin regulates MUC2 expression via B3GNT7. **(A)** Venn diagram was plotted to show the intersected genes from four groups, NS (OXTR^fl/fl^); KO (OXTR^△IEC^), 10 common genes were screened out. **(B)** The KEGG pathways for the Up-regulated mRNAs and down-regulated mRNAs. **(C and D)** Fold change (log2) in gene expression of in the IECs from OXTR^fl/fl^ (n=3) and OXTR^△IEC^ (n=3) mice treated with or without DSS, plotted against significance (−log10[p value]). Up-regulated genes (p-value < 0.05) are represented in red and down-regulated genes are labeled in blue. **(E)** RT-qPCR analysis of the expression of B3GNT7 in the IECs from OXTR^fl/fl^ and OXTR^△IEC^ mice. (n=5). **(F)** Western blot analysis of the expression of B3GNT7 in the IECs from OXTR^fl/fl^ and OXTR^△IEC^ mice. (n=5). **(G)** RT-qPCR analysis of the expression of B3GNT7 in the IECs from OXTR^fl/fl^ and OXTR^△IEC^ mice treated with 2.5%DSS. (n=3) **(H)** Western blot analysis of the expression of B3GNT7 in the IECs from OXTR^fl/fl^ and OXTR^△IEC^ mice treated with 2.5%DSS. (n=3). **(I and J)** Western blot analysis of the MUC2 level in LS174T cells **(I)** and colonic organoids **(J)** treated with OXT (1mmol) or siRNA (50 nmol). (n=5). Data are presented as the mean ± SEM. *P < 0.5. P-values were determined by using an unpaired two-tailed Student’s t-test.

### OXT/OXTR Induces MUC2 α1-3-Fucosylation for Protection Against Colitis

B3GNT7, which is involved in the combination of poly-N-acetyllactosamine chains and triggers α1-3-fucosylation of glycoproteins in IECs (20), has been exhibited to inhibit the metastasis of colon cancer cells (21). Since OXT facilitates the expression of B3GNT7 (Figure. 4), we explored whether OXT also promotes fucosylation in IECs. We evaluated the overall fucosylation status of glycoproteins using fucose-recognizing lectins in IEC lysates. The results showed attenuated binding of Lotus tetragonolobus lectin (LTL), which characteristically identifies α1-3-linked fucose, in OXTR-deficient IECs (Figure. 5A). Additionally, we compared fucosylation patterns in the colon tissues of OXTR^fl/fl^ and OXTR^△IEC^ mice using LTL staining and found reduced lectin binding in OXTR^△IEC^ mice (Figure. 5B). To understand the mechanisms underlying OXT-induced increased α1-3-fucosylation, we examined the expression levels of fucosyltransferases (FUTs) (22), B3GNTs, and glycosidases in IECs (23). Interestingly, except for B3gnt7, the transcription levels of other enzymes remained unchanged (Figure. S4, A-C). Furthermore, we observed reduced binding of Lycopersicon esculentum lectin (LEL), which recognizes GlcNAc residues in chitin and polyLacNAc chains,(20) in the colon tissues of OXTR^△IEC^ mice (Figure. 5C).

**Figure. 5.**
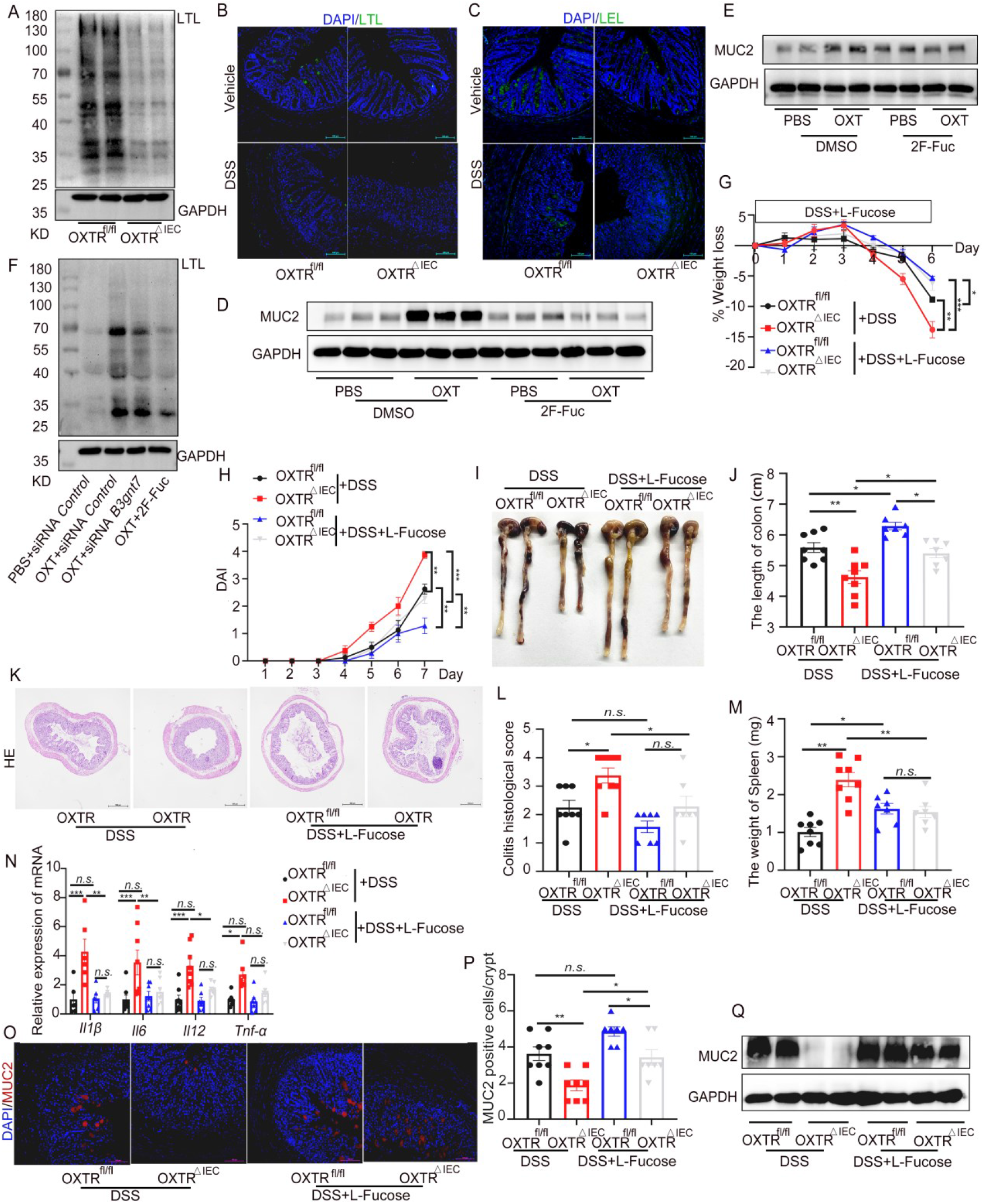
Fucosylation is responsible for oxytocin mediated protective effect in DSS-induced acute colitis. **(A)** Western blot analysis of the LTL level in the IECs from OXTR^fl/fl^ and OXTR^△IEC^ mice. (n=3) **(B)** Immunofluorescence staining of LTL (green) and nuclei (DAPI, blue), scale bar, 100 μm. (n=5) **(C)** Immunofluorescence staining of LEL (green) and nuclei (DAPI, blue), scale bar, 100 μm. (n=5) **(D and E)** Western blot analysis of the MUC2 level in LS174T cells **(D)** and colonic organoids **(E)** treated with OXT (1mmol) or 2F-Fuc (200 μmol). (n=5). **(F)** Western blot analysis of the LTL level in LS174T cells treated with siRNA, OXT, or 2F-Fuc. (n=5). G-Q) OXTR^fl/fl^ and OXTR^△IEC^ mice were treated with 2.5% DSS (n=8) or 200 mM L-Fucose (n=7). **(G)** Body weights were measured daily and are depicted as a percentage of initial body weight. **(H)** DAI score. Representative photos of the colons. (J) Statistical analysis of colon length. (K) Representative H&E-stained images.Scale bar, 500 μm. **(L)** Histologic injury score. **(M)** Statistical analysis of spleen weight. **(N)** RT-qPCR analysis of the relative expression of inflammatory factors in colon tissues. **(O)** Immunofluorescence staining of Muc2 (red) and nuclei (DAPI, blue), scale bar, 100 μm. **(P)** The number of MUC2-positive cells were counted randomly in crypts for each mouse. **(Q)** Western blot analysis of the MUC2 level in the IECs from OXTR^fl/fl^ and OXTR^△IEC^ mice treated with or without L-Fucose. Data are presented as the mean ± SEM. n.s, no significance. *P < 0.5, **P < 0.1, ***P < 0.01. P-values were determined by using a two-way analysis of variance (ANOVA).

The expression and O-glycosylation alterations of MUC2 mucins have been implicated in colorectal cancer(24). To examine the effects of fucosylation regulated by OXT on LS174T cells and colonic organoids, we found that the upregulation of MUC2 by OXT was inhibited in the presence of 2-fluoroperacetyl-fucose (2F-Fuc), a metabolic inhibitor of fucosylation (25), indicating that increased MUC2 expression depends on fucosylated structures (Figure. 5, D and E). Moreover, OXT can up-regulate the level of fucosylation, and this effect can be inhibited when B3GNT7 expression is reduced, which further indicates that OXT can regulate fucosylation through B3GNT7(Figure. 5F). To further investigate the role of fucosylation-mediated MUC2 in the maintenance of colitis, we tested the effect of L-Fucose supplementation, an essential substrate of fucosylation (26), in a DSS-induced colitis model. Remarkably, OXTR^△IEC^ mice treated with L-Fucose showed improved weight loss and disease activity index compared to those without L-Fucose (Figure. 5, G and H). Additionally, DSS-induced colon shortening was significantly ameliorated in OXTR^fl/fl^ and OXTR^△IEC^ mice receiving L-Fucose (Figure. 5, I and J). Histological damage and the levels of inflammation were also reduced in L-Fucose-treated mice, with less epithelial damage, crypt loss, spleen enlargement, and inflammatory factor content (Figure. 5, K-N). Importantly, the treatment of OXTR-deficient mice with L-Fucose led to a significant increase in colonic MUC2 levels (Figure. 5, O-Q). These findings suggest that L-Fucose treatment restores colonic MUC2 concentration in OXTR^△IEC^ mice and alleviates DSS-induced colitis.

### Treatment with OXT suppresses CAC development

Having established that mice with OXTR deficiency in intestinal epithelium exhibit more advanced colitis and colon cancer (Figure. 1-5), we further investigated the role of the OXT system in CAC through OXT supplementation (Figure. 6A). Mice treated with OXT exhibited longer colon lengths compared to control mice (Figure. 6, B and C). Furthermore, OXT-treated mice developed significantly fewer and smaller tumors in the colon compared to OXTR^fl/fl^ mice (Figure. 6, D and E). Statistical analysis confirmed reduced spleen size and suppressed inflammation levels in OXTR^fl/fl^ mice treated with OXT (Figure. 6, F and G). Histologically, OXTR^fl/fl^ mice treated with OXT showed reduced tumor areas, less colon injury, and immune cell infiltration (Figure. 6H). Moreover, the survival time was significantly prolonged in OXT-treated mice (Figure. 6I). These findings underscore the protective role of OXT in colitis-associated colon cancer.

**Figure. 6.**
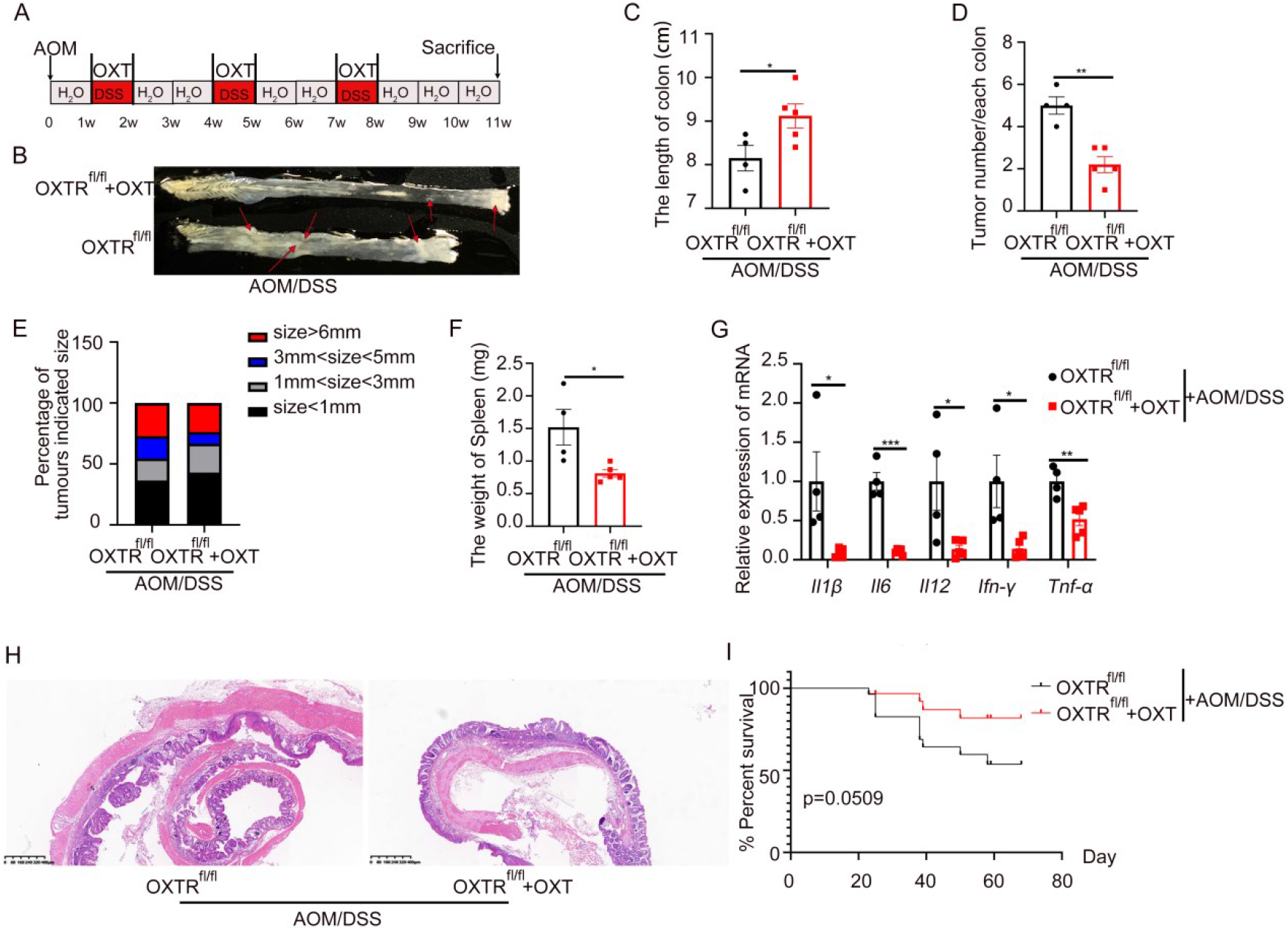
Oxytocin treatment reduces colon tumor burden. OXTR^fl/fl^ mice were treated with AOM and 3 cycles of 2% DSS (n=4) or DSS + OXT (n=5) to induce inflammation-driven colorectal cancer. **(A)** Flow chart of the induction procedure. **(B)** Representative photos of the colons. **(C)** Colon length. **(D)** Number of tumors in the colon. **(E)** The percentage of tumors by different diameters. **(F)** Statistical analysis of spleen weight. **(G)** Relative inflammatory cytokine mRNA levels in the colon were measured by real-time polymerase chain reaction. **(H)** Representative H&E-stained colon sections. **(I)** Survival curve assessed by log-rank (Mantel-Cox) test. Data are presented as the mean ± SEM. *P < 0.5, **P < 0.1, ***P < 0.01. P-values were determined by using C, D, F) unpaired two-tailed Student’s t-test or G) two-way analysis of variance (ANOVA).

### Decreased Expression of B3GNT7 Correlates with OXTR in Human Colitis

To understand the relevance of the decreased expression of OXTR and B3GNT7 in humans, we analyzed an independent cohort of colitis patients and found decreased expression of OXTR in colitis (Figure. 7A). Using patient samples, we observed reduced expression of OXTR in the colon tissues of UC patients compared to normal controls (Figure. 7, B and C). Consistent with the animal results, B3GNT7 (Figure. 7, D and E) and MUC2 (Figure. 7, E and F) expression was significantly lower and the expression level of FUTs (Figure. S4D) was unchanged in human colitis. Furthermore, we examined colon tissue and IECs from colitis-associated cancer patients (Figure. 7G) and an AOM/DSS mouse model (Figure. 7, H and I). We found reduced MUC2 expression in colitis-associated cancer. Additionally, in human UC patients and colitis-associated cancer patients, the expression of B3GNT7 was decreased and positively correlated with OXTR (Figure. 7, J-L). These results indicate that the downregulation of B3GNT7 is strongly associated with OXTR in human inflammation and colonic carcinogenesis.

**Figure. 7.**
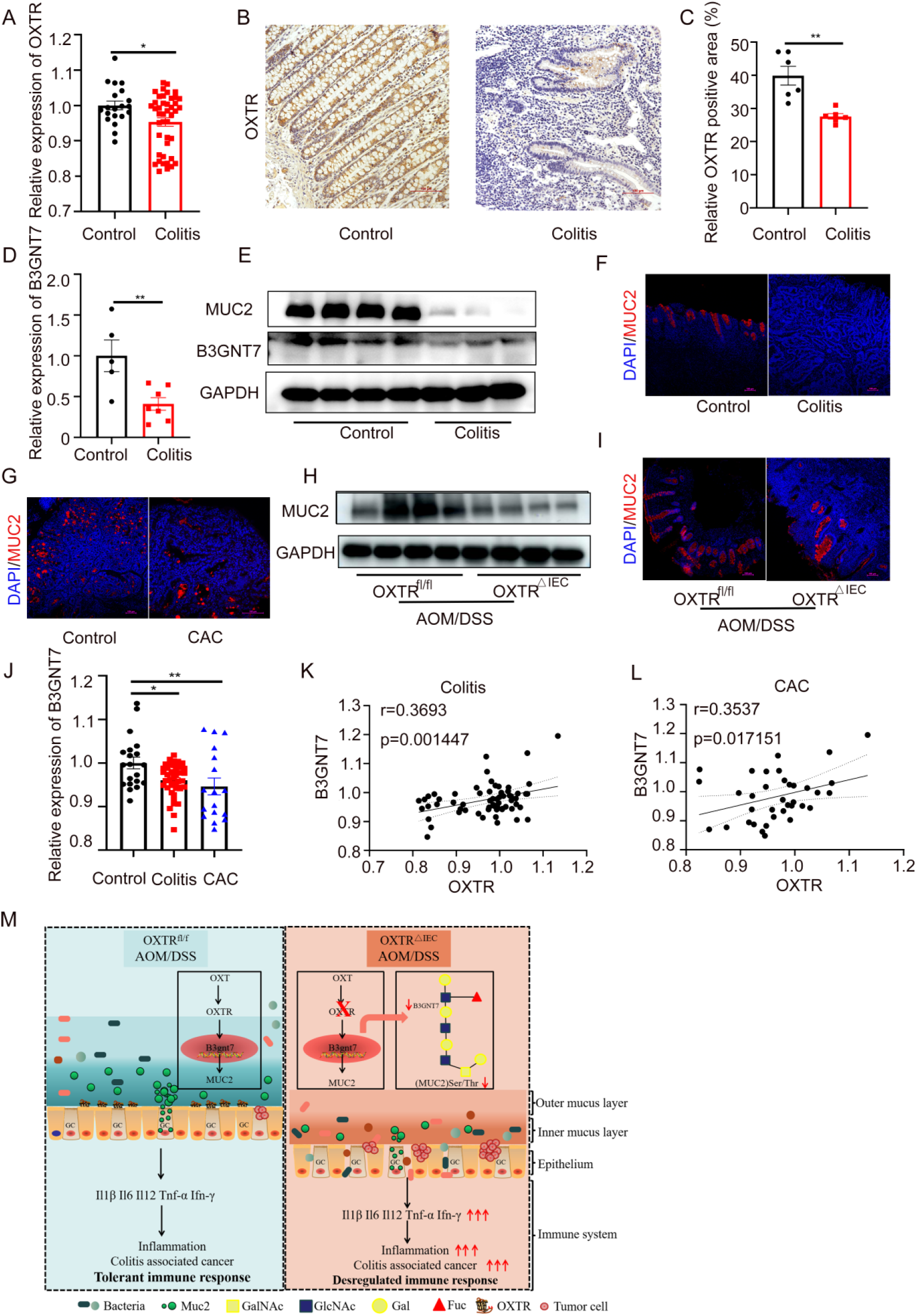
OXTR and B3GNT7 expressions in human colitis. **(A)** OXTR gene expression in human normal and colitis tissue. (GSE37283 and GSE47908). **(B)** Representative images of OXTR staining on human normal and colitis tissue. Scale bar, 100 μm. **(C)** The quantifications of OXTR-positive area. (n=6). **(D)** RT-qPCR analysis of the relative expression of B3GNT7 in human normal (n=5) and colitis (n=7) tissue. **(E)** Western blot analysis of the MUC2 and B3GNT7 level in human normal (n=4) and colitis tissue (n=3). **(F)** Immunofluorescence staining of Muc2 (red) and nuclei (DAPI, blue) in human normal (n=5) and colitis tissue (n=5). Scale bar, 100 μm. **(G)** Immunofluorescence staining of Muc2 (red) and nuclei (DAPI, blue) in human normal (n=3) and CAC tissue (n=3). Scale bar, 100 μm. **(H and I)** Western blot analysis **(H)** and immunofluorescence staining **(I)** of MUC2 in OXTR^fl/fl^ and OXTR^△IEC^ mice treated with AOM/DSS. (n=5). (J) B3GNT7 gene expression in human normal, colitis and CAC tissue. (GSE37283 and GSE47908). **(K and L)** Correlations between the expression of B3GNT7 and OXTR in human colitis **(K)** and CAC **(L). (M)** Schematic model depicting the functions of intestinal OXTR in CAC. Data are presented as the mean ± SEM. *P < 0.5, **P < 0.1. P-values were determined by using A, C, D) unpaired two-tailed Student’s t-test or J) two-way analysis of variance (ANOVA).

## Discussion

In this study, we present compelling evidence supporting the involvement of oxytocin in the formation of colonic mucus and its protective role in colitis and CAC tumorigenesis. We demonstrate that the depletion of OXTR specifically in intestinal epithelial cells contributes to the development of experimental colitis induced by DSS and CAC induced by a combination of DSS and azoxymethane. To unravel the underlying molecular mechanisms, we elucidate that oxytocin orchestrates the maturation of colonic mucus proteins through B3GNT7-mediated fucosylation. Our study uncovers a novel regulatory pathway wherein oxytocin controls the fucosylation process crucial for the formation of colonic mucin. Remarkably, treatment with oxytocin not only enhances mucin levels but also ameliorates tumor burden in CAC development (Figure. 7M).

While oxytocin is widely recognized for its involvement in behaviors such as social bonding, reproduction, childbirth, and postpartum functions (27), we extend the understanding of its role in the gastrointestinal system. Enteric nerves can release oxytocin, and various cells in the intestine express OXTR (28), indicating its multifaceted functions beyond the traditional domains. Recently, a study showed that stimulation of oxytocin neurons inhibits CAC progression,(10) which implies that oxytocin may be involved in CAC, however, the mechanism is unclear. Here, our findings provide an explanation that activation of oxytocin signaling in the colonic epithelium promoted protective mucin protein MUC2 maturation by B3GNT7-mediated fucosylation. MUC2 is the first defense layer of the colon and plays a vital role in colitis and CAC development. Hormones can regulate plenty of gene expression through the activation of transcription factors, the oxytocin-involved post-translational modification of MUC2 we identified may be a general way for hormones’ physiology functions.

Interestingly, prior research utilizing systematic bioinformatics analysis has suggested that high levels of OXTR mRNA are associated with a poor prognosis in colon cancer, implicating OXTR as a potential marker for human colon cancer (29). However, our data unexpectedly reveal a protective role of OXTR in both DSS-induced colitis and AOM/DSS-induced CAC tumorigenesis mouse models. Moreover, our analysis of human colitis and CAC samples reveals downregulation of OXTR expression, indicating complex and context-dependent roles of oxytocin signaling in these conditions. Notably, we hypothesize that neoplastic lesions in colitis, unlike sporadic colon cancer, may largely operate independently of Wnt signaling (30), warranting further investigation into the specific function of oxytocin signaling in Wnt-driven colon cancer.

Fucosylation has been established as a crucial modification for maintaining normal colon homeostasis (31), and previous studies have highlighted the protective effects of fucose supplementation through FUT2-mediated fucosylation against colitis onset (32). In our study, we uncover an alternative mechanism involving B3GNT7-mediated fucosylation, which contributes to colonic mucin formation and is regulated by the intricate gut-brain axis. Remarkably, we find that in the gut, mucin production is determined by FUT8 rather than FUT2 (33). FUT8, as the sole enzyme responsible for α1-6–linked fucosylation, adds fucose to the innermost GlcNAc residue of an N-linked glycan (34). While oxytocin does not directly affect the mRNA levels of fucosyltransferases, our findings suggest that oxytocin is involved in α1-3–linked fucosylation through the non-fucosyltransferase B3GNT7. Hence, we propose that B3GNT7-mediated α1-3–linked fucosylation plays a crucial role in the maturation of MUC2 protein, while FUT8-mediated α1-6–linked fucosylation governs the secretion of MUC2 in the colon. Both these processes are essential for protecting the digestive tract from pathogens and maintaining intestinal health.

In summary, our study provides comprehensive evidence highlighting the crucial role of oxytocin in maintaining colon homeostasis and protecting against colitis and colorectal cancer (CAC) tumorigenesis. We elucidate the involvement of oxytocin in regulating intestinal mucin production through the fucosylation pathway mediated by B3GNT7. Intriguingly, reduced expression of OXTR and B3GNT7 is observed in patients with colitis and CAC, underscoring the clinical relevance of our findings. Considering the FDA approval and good tolerability of oxytocin in humans, it represents a promising and cost-effective therapeutic avenue for patients suffering from colitis and CAC.

## Materials and Methods

### Experimental Animal Studies

OXTR^flox/flox^ (OXTR^fl/fl^) mice and villin-cre mice (B6.Cg-Tg[Vil1-cre]997Gum/J, 004586) in the C57BL/6 genetic background were obtained from Jackson Laboratory. Intestine-specific OXTR null mice (OXTR^△IEC^) were generated by cross-fertilizing the transgenic mice. Sex-and age-matched littermate mice were used for all experiments. The mice were bred and maintained in specific pathogen-free conditions at the animal facility of the School of Basic Medical Sciences. All animal experiments were conducted following the guidelines of the National Institutes of Health Guide for the Care and Use of Laboratory Animals and were approved by the Scientific Investigation Board of Shandong University (#ECSBMSSDU2019-2-048), Jinan, Shandong Province, China.

### CAC and Colitis Model

To induce the CAC model, 8-week-old mice were intraperitoneally injected with (10 mg/kg) AOM (Sigma) (18). After 7 days, 2.2% DSS (MP Biomedicals) was administered in drinking water for 5 days, followed by regular water for 14 days. This 5-day cycle of 2.2% DSS was repeated twice, and mice were sacrificed after the last DSS cycle. For the AOM alone–induced colon cancer model, 8-week-old mice were intraperitoneally injected with AOM (10 mg/kg) once a week for 6 weeks, followed by a five-month waiting period to induce spontaneous colonic carcinogenesis. Mice were euthanized by CO2 asphyxiation. For the chronic DSS model, 8-week-old mice were administered 2.2% DSS in autoclaved drinking water for 6 days, followed by regular water for 5 days. This 5-day cycle of 2.2% DSS was repeated twice, and mice were sacrificed after the last DSS cycle. For the acute DSS model, 8-week-old mice were administered 2.5% DSS in autoclaved drinking water for 6 days.

### Fucose Supplementation

To investigate the effect of L-fucose, (200 mM) L-Fucose (Solarbio) dissolved in saline, was given to mice during the DSS administration period (35).

### Pharmacological Experiment

LS174T cells were grown in supplemented DMEM for three days, while organoids were cultured for 7 days in a complete medium to achieve differentiated goblet cells. Cells were then placed in serum-free media for 16 to 18 hours before treatment with inhibitors, including (200 μM) 2F-Fuc or 1 μmol L-368,899, or vehicle control for 30 minutes. Cells were then stimulated with OXT at the indicated time and concentrations.

### Transfection of siRNA

Cells were transfected with B3gnt7 siRNAs or control siRNAs using INTERFERin (LS174T) or lip3000 (crypt) according to the manufacturer’s instructions. The targeted siRNA sequences were designed and synthesized by Generay Biotechnology (Shanghai).

### Analysis of Human Patient Datasets and Human Cohort

Gene expression data correlation analysis in colonic mucosal biopsies from human colitis patients and cancer patients was obtained from Gene Expression Omnibus (GEO) databases GSE37283, GSE1113002, and GSE47908. Colonic tissue biopsies were obtained from Qilu Hospital of Shandong University, and informed consent was obtained from all patients. The use of human tissues was approved by the Medical Institutional Ethics Committee of Qilu Hospital, Shandong University, China (#KYLL-202210-049).

### Statistical Analysis

The statistical significance of group differences was assessed using GraphPad Prism 8.0 (GraphPad Software). Data are presented as the mean ± SEM and based on experiments performed at least in triplicate. The two-way ANOVA test was used for the comparison between the two samples, and the Unpaired Student’s t-test was used for the comparison between the two groups. A p-value less than 0.05 was considered statistically significant.

### Other detailed information for the materials and methods is provided in the supplemental materials

## Acknowledgments

Special thanks go to Prof. Zhen Junhui from the Pathology Department of Qilu Hospital for providing UC and CAC samples.

## Authors’ contributions

W.X. and D.C performed research, analyzed the data, and wrote the manuscript; M.G and Y.N performed research and analyzed the data; J. G and D. Z performed research; J. G and X. X provided materials; L. L, L. L, and Y. L were responsible for the design of the study, interpretation of data and writing of the manuscript; W.X, X. Z and J. L conceptualized the project and funded the work.

## Data and materials availability

All data needed to evaluate the conclusions in the paper are present in the paper and/or the Supplementary Materials. The RNA-seq data were uploaded to Gene Expression Omnibus (GEO) with the accession code PRJNA996718. The GEO accession codes for previously published datasets used in this study are GSE37283, GSE1113002, and GSE47908.

